## Supplementary figures and tables for "NetTCR 2.2 - Improved TCR specificity predictions by combining pan- and peptide-specific training strategies, loss-scaling and integration of sequence similarity"

**Supplementary material**

| **Peptide** | **Pre Reduction Count** | **Removed in first reduction** | **Removed in second reduction** | **Post Reduction Count** |
| --- | --- | --- | --- | --- |
| GILGFVFTL | 1897 | 645 | 127 | 1125 |
| RAKFKQLL | 1065 | 114 | 17 | 934 |
| KLGGALQAK | 912 | 8 | 2 | 902 |
| AVFDRKSDAK | 725 | 5 | 4 | 716 |
| ELAGIGILTV | 435 | 6 | 3 | 426 |
| NLVPMVATV | 384 | 43 | 11 | 330 |
| IVTDFSVIK | 323 | 13 | 2 | 308 |
| LLWNGPMAV | 322 | 72 | 21 | 229 |
| CINGVCWTV | 231 | 4 | 1 | 226 |
| GLCTLVAML | 278 | 59 | 7 | 212 |
| SPRWYFYYL | 158 | 4 | 5 | 149 |
| ATDALMTGF | 128 | 21 | 4 | 103 |
| DATYQRTRALVR | 100 | 4 | 3 | 93 |
| KSKRTPMGF | 115 | 14 | 12 | 89 |
| YLQPRTFLL | 69 | 6 | 1 | 62 |
| HPVTKYIM | 60 | 5 | 2 | 53 |
| RFPLTFGWCF | 58 | 7 | 0 | 51 |
| GPRLGVRAT | 51 | 3 | 0 | 48 |
| CTELKLSDY | 48 | 0 | 0 | 48 |
| RLRAEAQVK | 47 | 0 | 0 | 47 |
| RLPGVLPRA | 43 | 0 | 0 | 43 |
| SLFNTVATLY | 38 | 0 | 0 | 38 |
| RPPIFIRRL | 40 | 2 | 2 | 36 |
| VLFGLGFAI | 32 | 1 | 0 | 31 |
| FEDLRLLSF | 31 | 0 | 0 | 31 |
| FEDLRVLSF | 36 | 0 | 13 | 23 |

***Supplementary Table 1*** *– Overview of number of observations removed as a result of the Hobohm 1 redundancy reductions at a 95% similarity threshold. The first reduction took care of redundancy of TCRs within each peptide separately, whereas the second reduction took care of redundancy between TCRs of different peptides.*

| **Peptide Sequence** | **Organism** | **HLA** | **# positive TCRs** |
| --- | --- | --- | --- |
| GILGFVFTL | Influenza A virus | HLA-A*02:01 | 1125 |
| RAKFKQLL | Epstein Barr virus | HLA-B*08:01 | 934 |
| KLGGALQAK | Human CMV | HLA-A*03:01 | 902 |
| AVFDRKSDAK | Epstein Barr virus | HLA-A*11:01 | 716 |
| ELAGIGILTV | Melanoma neoantigen | HLA-A*02:01 | 426 |
| NLVPMVATV | Human CMV | HLA-A*02:01 | 330 |
| IVTDFSVIK | Epstein Barr virus | HLA-A*11:01 | 308 |
| LLWNGPMAV | Yellow fever virus | HLA-A*02:01 | 229 |
| CINGVCWTV | Hepatitis C virus | HLA-A*02:01 | 226 |
| GLCTLVAML | Epstein Barr virus | HLA-A*02:01 | 212 |
| SPRWYFYYL | SARS-CoV2 | HLA-B*07:02 | 149 |
| ATDALMTGF | Hepatitis C virus | HLA-A*01:01 | 103 |
| DATYQRTRALVR | Influenza A virus | HLA-A*68:01 | 93 |
| KSKRTPMGF | Hepatitis C virus | HLA-B*57:01 | 89 |
| YLQPRTFLL | SARS-CoV2 | HLA-A*02:01 | 62 |
| HPVTKYIM | Hepatitis C virus | HLA-B*08:01 | 53 |
| RFPLTFGWCF | HIV-1 | HLA-A*24:02 | 51 |
| CTELKLSDY | Influenza A virus | HLA-A*01:01 | 48 |
| GPRLGVRAT | Hepatitis C virus | HLA-B*07:02 | 48 |
| RLRAEAQVK | Epstein Barr virus | HLA-A*03:01 | 47 |
| RLPGVLPRA | AML neoantigen | HLA-A*02:01 | 43 |
| SLFNTVATLY | HIV-1 | HLA-A*02:01 | 38 |
| RPPIFIRRL | Epstein Barr virus | HLA-B*07:02 | 36 |
| FEDLRLLSF | Influenza A virus | HLA-B*37:01 | 31 |
| VLFGLGFAI | T1D neoantigen | HLA-A*02:01 | 31 |
| FEDLRVLSF | Influenza A virus | HLA-B*37:01 | 23 |

***Supplementary Table 2*** *– Overview of source organism for each epitope, as well as the HLA allele which they bind to.*

| **Peptide Sequence** | **Not 10X** | **10X** | **Total** |
| --- | --- | --- | --- |
| GILGFVFTL | 426 | 699 | 1125 |
| RAKFKQLL | 0 | 934 | 934 |
| KLGGALQAK | 0 | 902 | 902 |
| AVFDRKSDAK | 0 | 716 | 716 |
| ELAGIGILTV | 55 | 371 | 426 |
| NLVPMVATV | 154 | 176 | 330 |
| IVTDFSVIK | 0 | 308 | 308 |
| LLWNGPMAV | 229 | 0 | 229 |
| CINGVCWTV | 75 | 151 | 226 |
| GLCTLVAML | 95 | 117 | 212 |
| SPRWYFYYL | 149 | 0 | 149 |
| ATDALMTGF | 0 | 103 | 103 |
| DATYQRTRALVR | 93 | 0 | 93 |
| KSKRTPMGF | 0 | 89 | 89 |
| YLQPRTFLL | 54 | 8 | 62 |
| HPVTKYIM | 0 | 53 | 53 |
| RFPLTFGWCF | 51 | 0 | 51 |
| GPRLGVRAT | 0 | 48 | 48 |
| CTELKLSDY | 48 | 0 | 48 |
| RLRAEAQVK | 0 | 47 | 47 |
| RLPGVLPRA | 0 | 43 | 43 |
| SLFNTVATLY | 0 | 38 | 38 |
| RPPIFIRRL | 24 | 12 | 36 |
| FEDLRLLSF | 31 | 0 | 31 |
| VLFGLGFAI | 31 | 0 | 31 |
| FEDLRVLSF | 23 | 0 | 23 |

***Supplementary Table 3*** *– Overview of the number of positive observations coming from 10X sequencing.*

| **Peptide** | **Pre Reduction Count** | **Post Reduction Count** | **Percent Redundant** |
| --- | --- | --- | --- |
| All | 2445 | 1960 | 19.8% |
| GILGFVFTL | 544 | 301 | 44.7% |
| NLVPMVATV | 274 | 242 | 11.7% |
| YLQPRTFLL | 267 | 227 | 15.0% |
| TTDPSFLGRY | 193 | 187 | 3.1% |
| LLWNGPMAV | 188 | 175 | 6.9% |
| CINGVCWTV | 183 | 179 | 2.2% |
| GLCTLVAML | 146 | 91 | 37.7% |
| ATDALMTGF | 104 | 78 | 25.0% |
| LTDEMIAQY | 100 | 94 | 6.0% |
| SPRWYFYYL | 92 | 92 | 0.0% |
| KSKRTPMGF | 85 | 63 | 25.9% |
| NQKLIANQF | 56 | 53 | 5.4% |
| HPVTKYIM | 48 | 41 | 14.6% |
| TPRVTGGGAM | 45 | 44 | 2.2% |
| NYNYLYRLF | 44 | 42 | 4.6% |
| GPRLGVRAT | 40 | 37 | 7.5% |
| RAQAPPPSW | 36 | 14 | 61.1% |

***Supplementary Table 4*** *– Overview of number of TCRs for each peptide in the IMMREP 2022 training dataset before and after redundancy reduction.*

| **Metric** | **PCC to Optimal Alpha** | **P-value** |
| --- | --- | --- |
| CNN AUC | -0.1101 | 0.2123 |
| TCRbase AUC | 0.3056 | 0.0004 |
| CNN AUC 0.1 | -0.0809 | 0.3602 |
| TCRbase AUC 0.1 | 0.2068 | 0.0183 |

***Supplementary Table 5*** *– Pearson Correlation Coefficients (PCC) between the optimal α scaling factor and performance in terms of AUC and AUC 0.1 of the pre-trained CNN model and TCRbase model, respectively, for the validation partitions. Each partition was considered as a separate sample. P-values for the null hypothesis that the performance and optimal α are uncorrelated are also shown.*

| **Peptide** | **Pre Reduction Count** | **Post Reduction Count** | **Percent Redundant** |
| --- | --- | --- | --- |
| All | 619 | 467 | 24.56% |
| GILGFVFTL | 136 | 58 | 57.35% |
| NLVPMVATV | 69 | 54 | 21.74% |
| YLQPRTFLL | 67 | 53 | 20.90% |
| TTDPSFLGRY | 49 | 47 | 4.08% |
| LLWNGPMAV | 47 | 44 | 6.38% |
| CINGVCWTV | 46 | 46 | 0.00% |
| GLCTLVAML | 37 | 23 | 37.84% |
| ATDALMTGF | 26 | 22 | 15.38% |
| LTDEMIAQY | 25 | 23 | 8.00% |
| SPRWYFYYL | 24 | 24 | 0.00% |
| KSKRTPMGF | 22 | 13 | 40.91% |
| NQKLIANQF | 15 | 15 | 0.00% |
| TPRVTGGGAM | 12 | 12 | 0.00% |
| HPVTKYIM | 12 | 10 | 16.67% |
| NYNYLYRLF | 12 | 9 | 25.00% |
| GPRLGVRAT | 11 | 11 | 0.00% |
| RAQAPPPSW | 9 | 3 | 66.67% |

***Supplementary Table 6*** *– Degree of redundancy between the IMMREP test and training data, when using a 95% kernel similarity threshold for redundancy within each peptide. The redundancy reduction was performed on both positive and negative observations. The counts and percentages, however, only refers to the positive observations.*

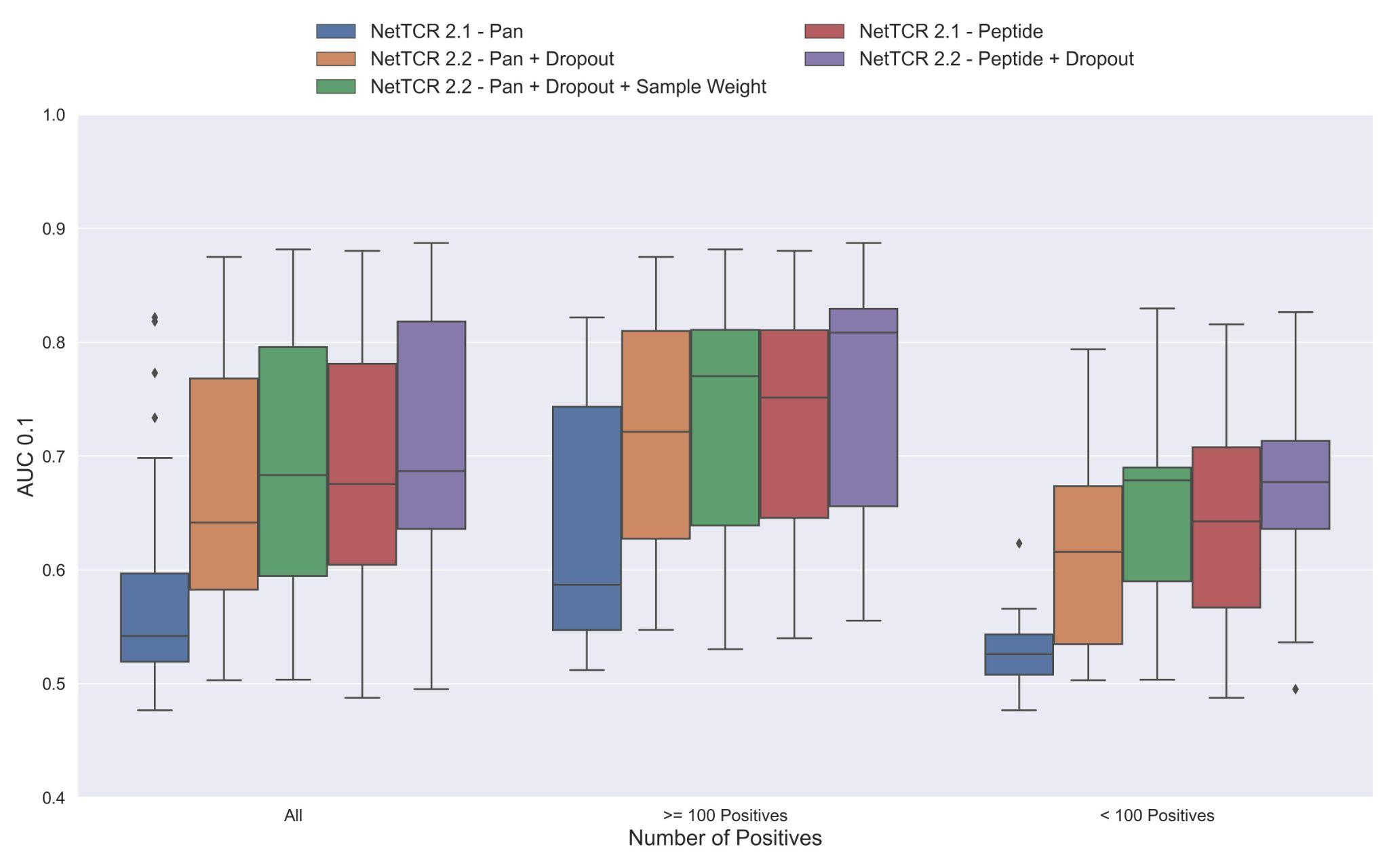
***Supplementary Figure 1*** *– Boxplot of AUC 0.1 of the pan- and peptide-specific NetTCR 2.1 and 2.2 models, respectively. The NetTCR 2.2 models include the updates to the model architecture, with the primary change being the introduction of dropout for the concatenated max-pooling layer (dropout rate = 0.6). Both the introduction of dropout and sample weights are shown to result in considerably improved performance for the pan-specific model. Separate boxplots are shown for all peptides, as well as separately for peptides with at least 100 positive observations and peptides with less than 100 positive observations, to highlight the effect of introducing dropout and sample weight for the least abundant peptides.*

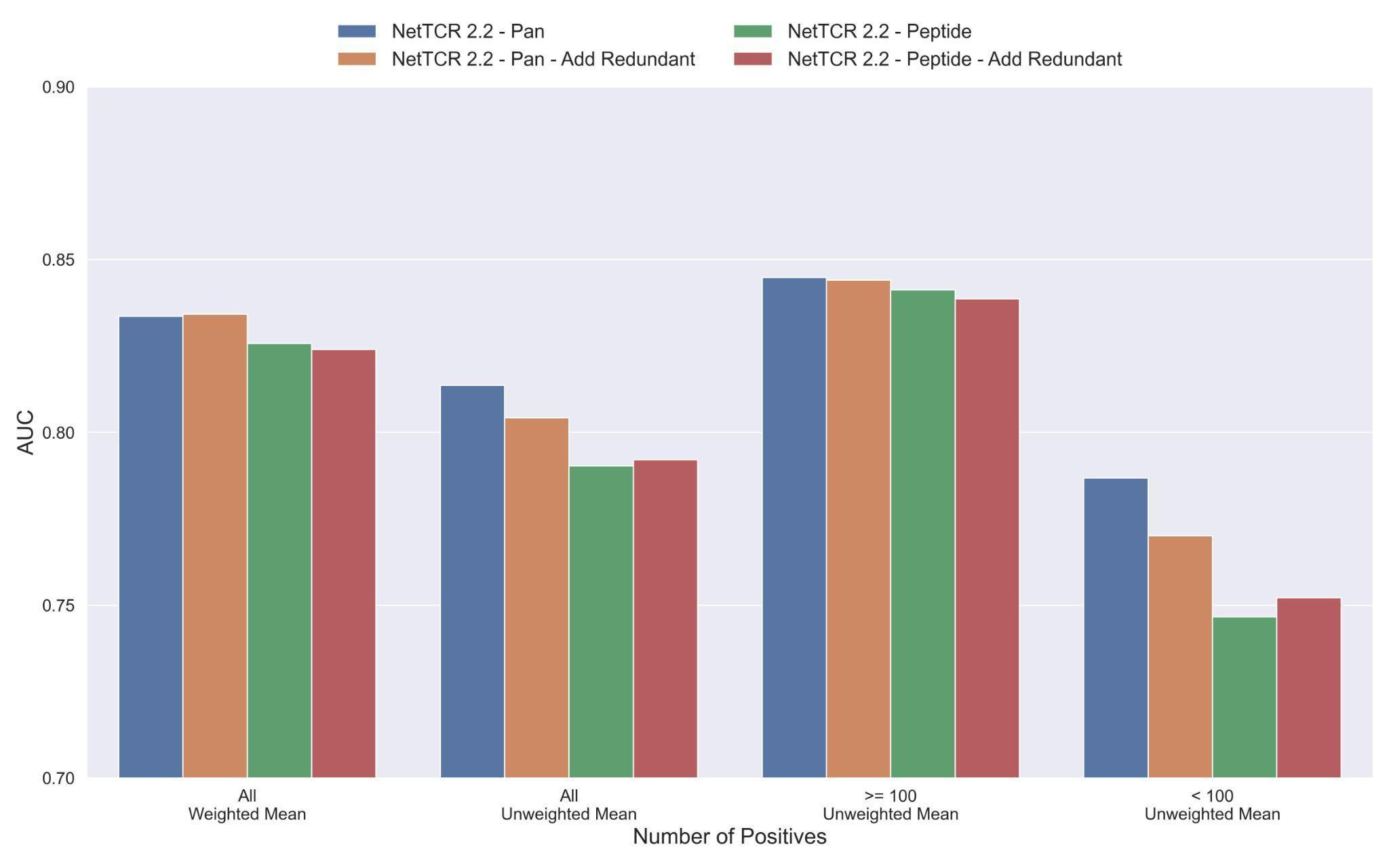
***Supplementary Figure 2 -*** *Mean AUC of the pan-specific and peptide-specific NetTCR 2.2 models, when training on the original redundancy reduced training data, and the dataset were redundant observations were added back to (“Add Redundant”). The AUC is reported in terms of weighted and unweighted mean across all peptides, as well as unweighted mean when the data is split into peptides with at least 100 positive observations, and less than 100 positive observations .*

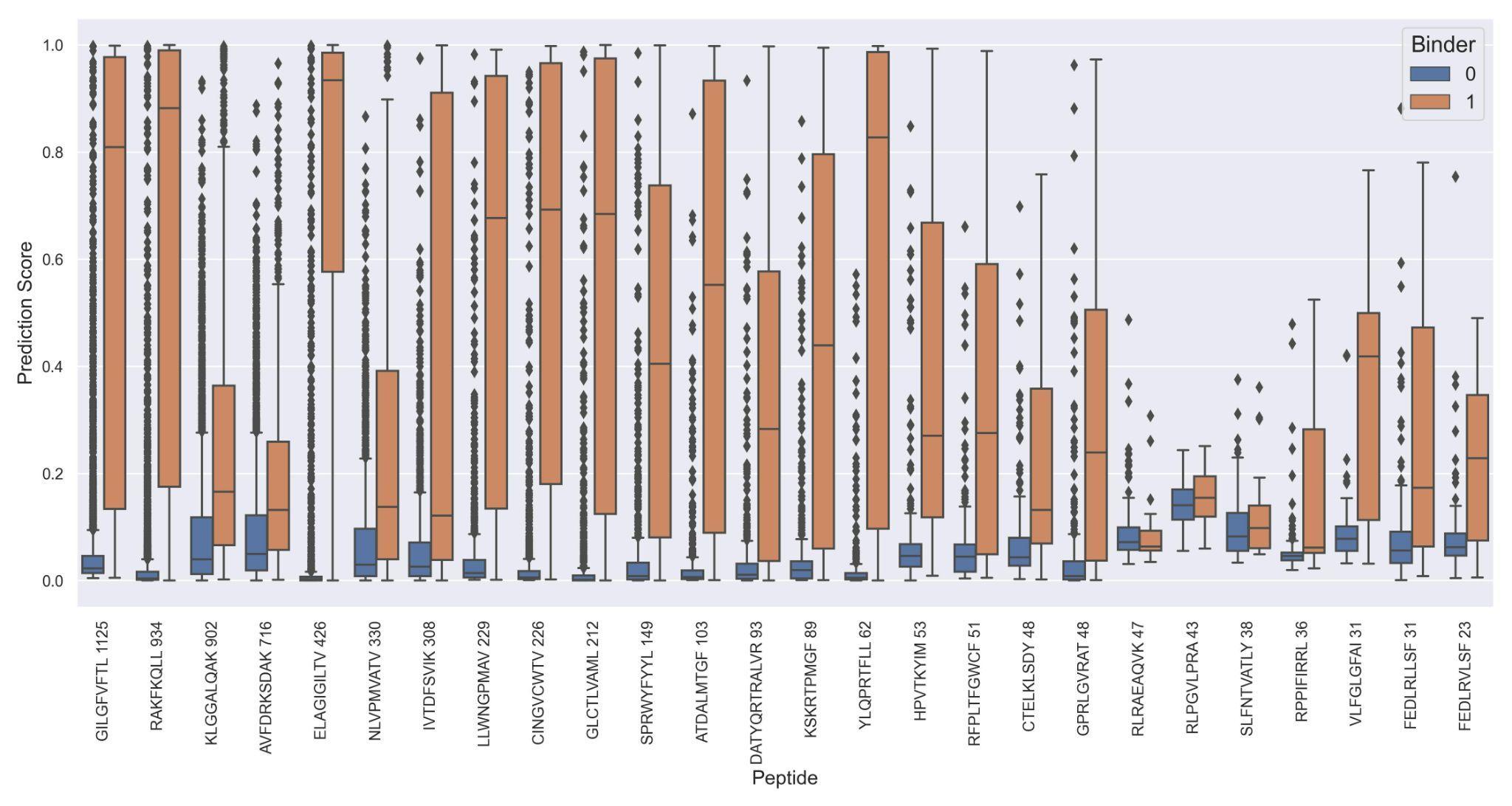

***Supplementary Figure 3*** *– Prediction values on the full test data for each peptide when predicted using the NetTCR 2.2 - Peptide model.*

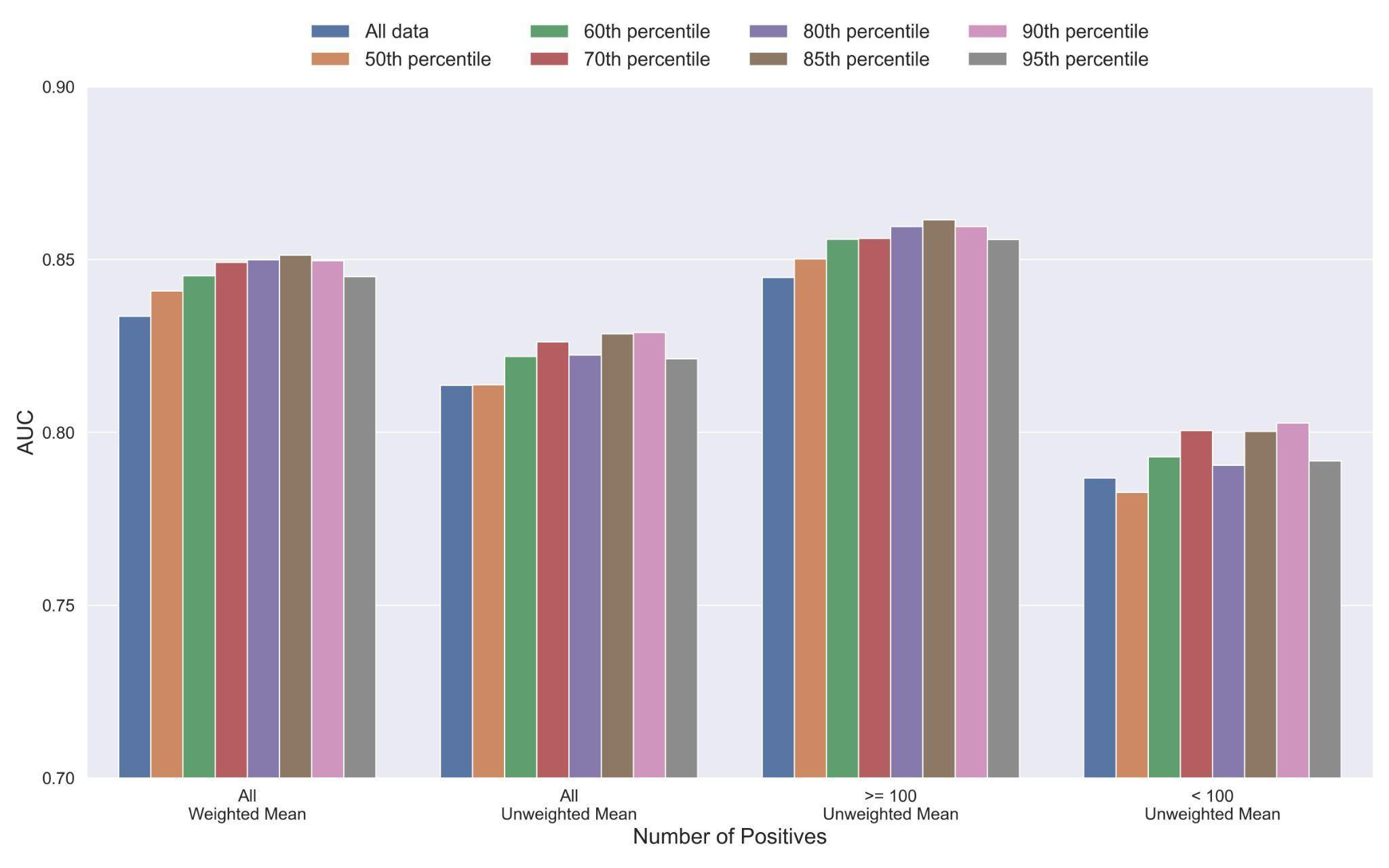

***Supplementary Figure 4*** *- Mean AUC of the pan-specific NetTCR 2.2 models when trained on datasets with potential outliers removed. The percentile refers to the threshold of prediction scores used for removing observations (see Material and Methods), and the higher the percentile is, the more observations are removed from training. The AUC is reported in terms of weighted and unweighted mean across all peptides, as well as unweighted mean when the data is split into peptides with at least 100 positive observations, and less than 100 positive observations .*

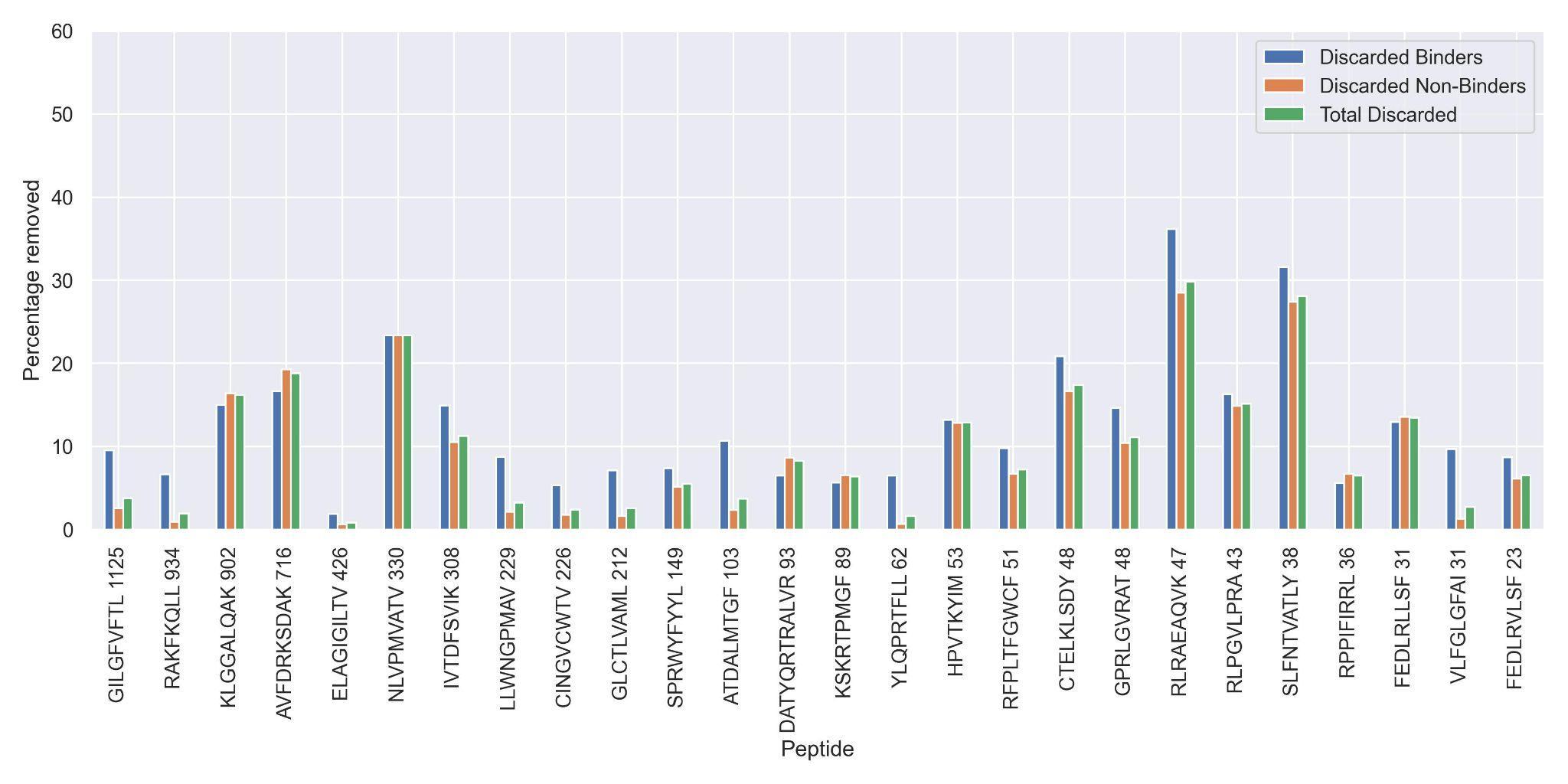

***Supplementary Figure 5*** *– Percentage of observations discarded for the 70^th^ percentile limited dataset, as a result of the removal of potential outliers.*

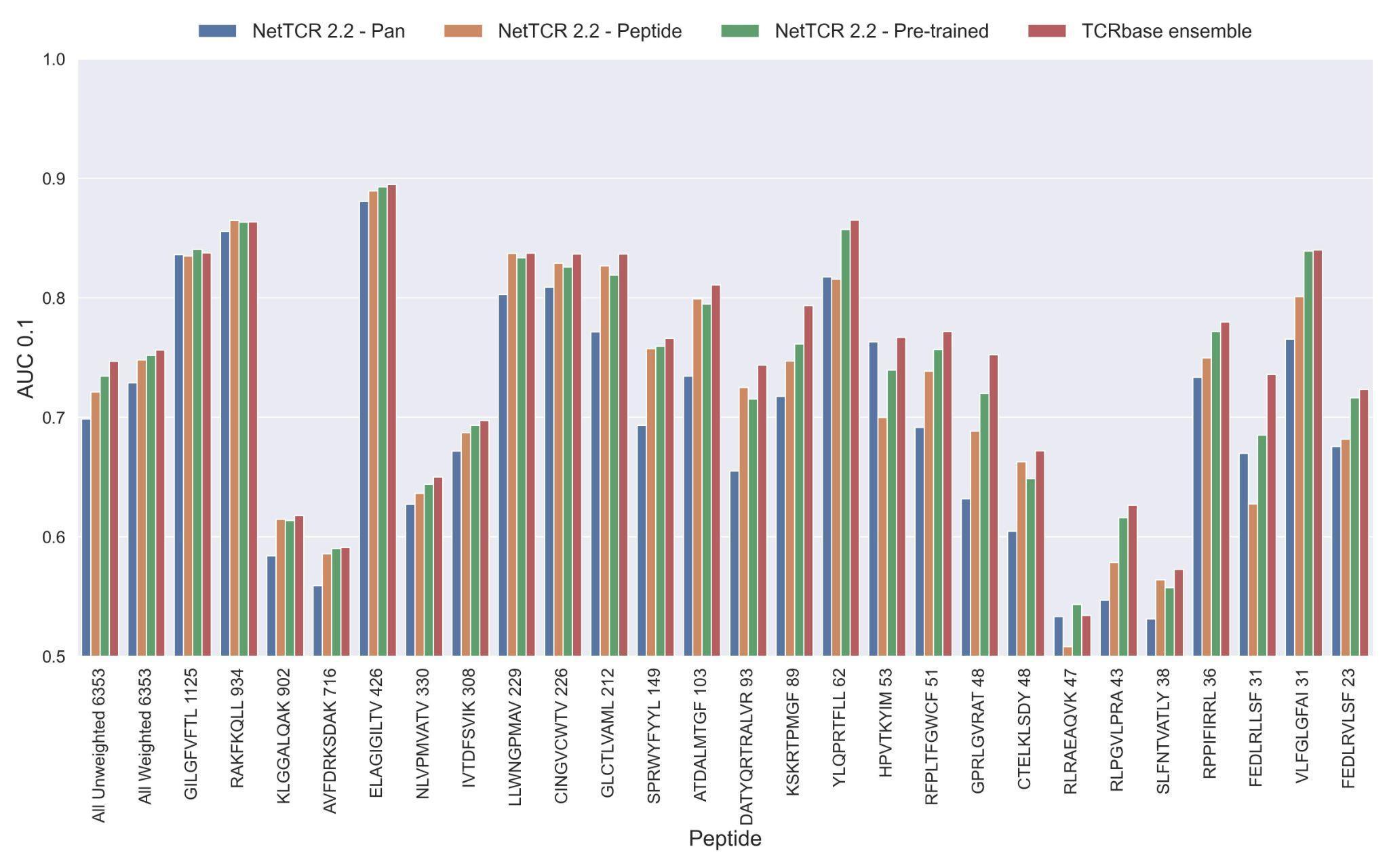

***Supplementary Figure 6* -** *Per peptide performance of the updated peptide-specific, pan-specific, and pre-trained CNN in terms of AUC 0.1, when trained on the limited training dataset and evaluated on the full dataset. The peptides are sorted based on the number of positive observations from most abundant to least abundant, with the number of positive observations listed next to the peptide sequence. The unweighted (direct) mean of AUC across all peptides is shown furthest to the left, while the weighted mean is shown second furthest to the left. The weighted mean is weighted by the number of positive observations per peptide and puts more emphasis on the peptides with the most observations.*

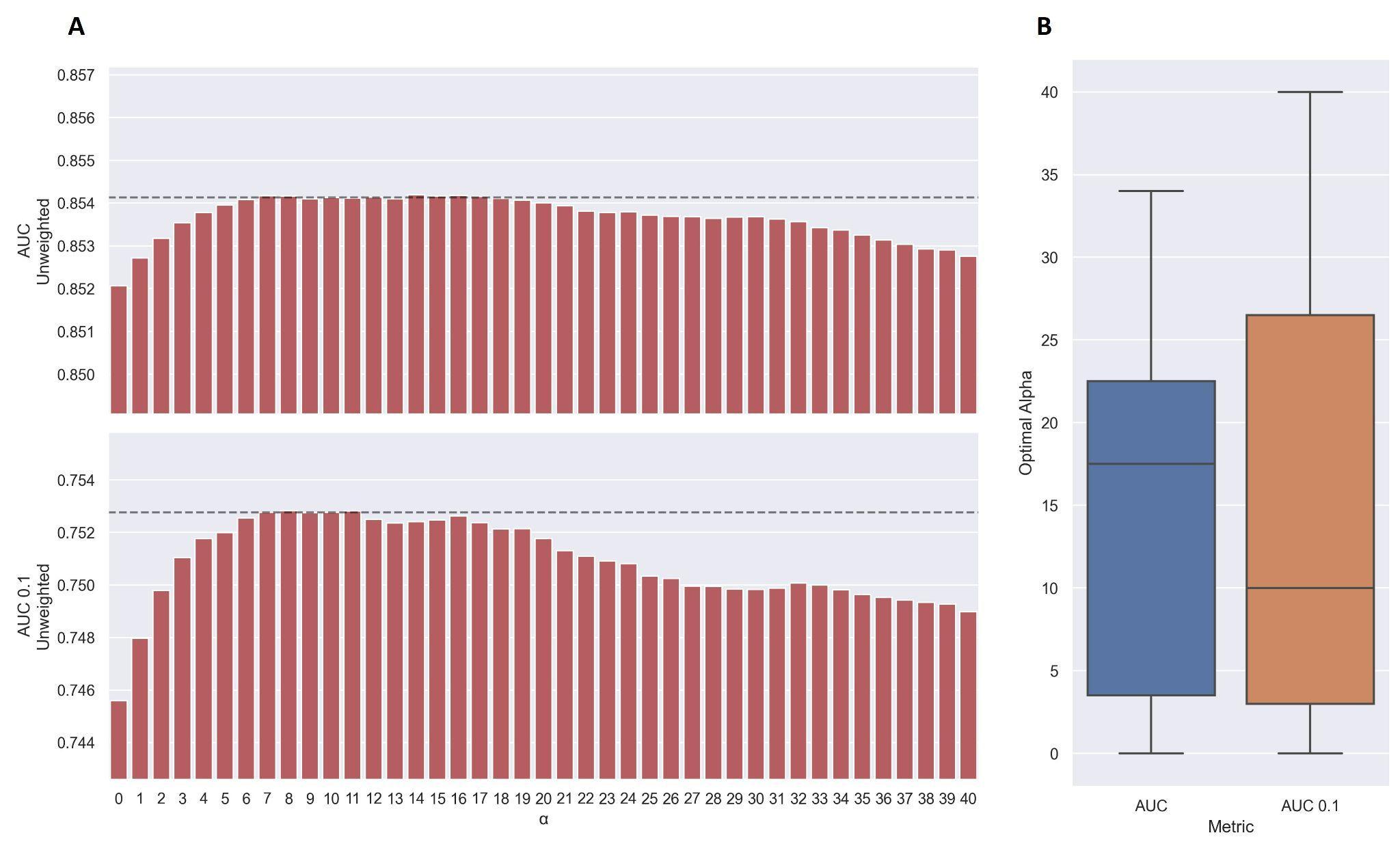

***Supplementary Figure 7*** *–* ***(A)*** *Performance of TCRbase ensemble as a function of α. The predictions of the pre-trained model ensemble (trained on the limited dataset) on the test partitions (full data) were scaled by the kernel similarity to known binders, as given by TCRbase with a weight of [1,1,3,1,1,3], to a power of α. The performance is given as the unweighted mean performance across all 26 peptides, in terms of AUC and AUC 0.1. The dashed line shows the performance when α is set to 10, which strikes a good balance between AUC and AUC 0.1. An α of zero corresponds to the model ensemble without the TCRbase scaling.* ***(B)*** *Boxplot of the optimal alpha scaling factor per cross-validation model, when evaluated on the validation partitions.*

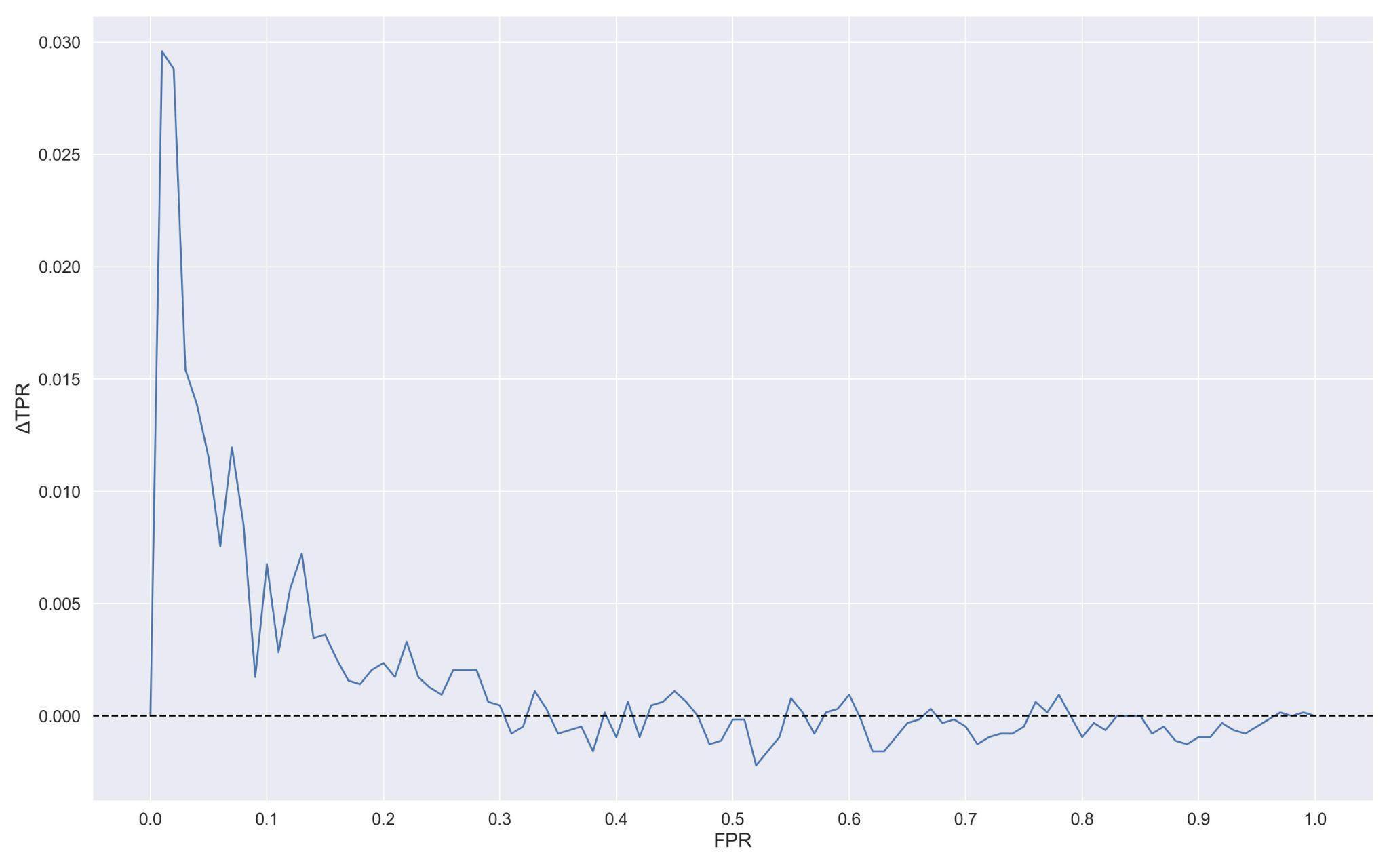

***Supplementary Figure 8*** *– Difference in true positive rate (TPR) between TCRbase ensemble (pre-trained + TCRbase models) and pre-trained models as a function of false positive rate (FPR). A positive ΔTPR corresponds to an increased performance of the TCRbase ensemble compared to the pre-trained models alone.*

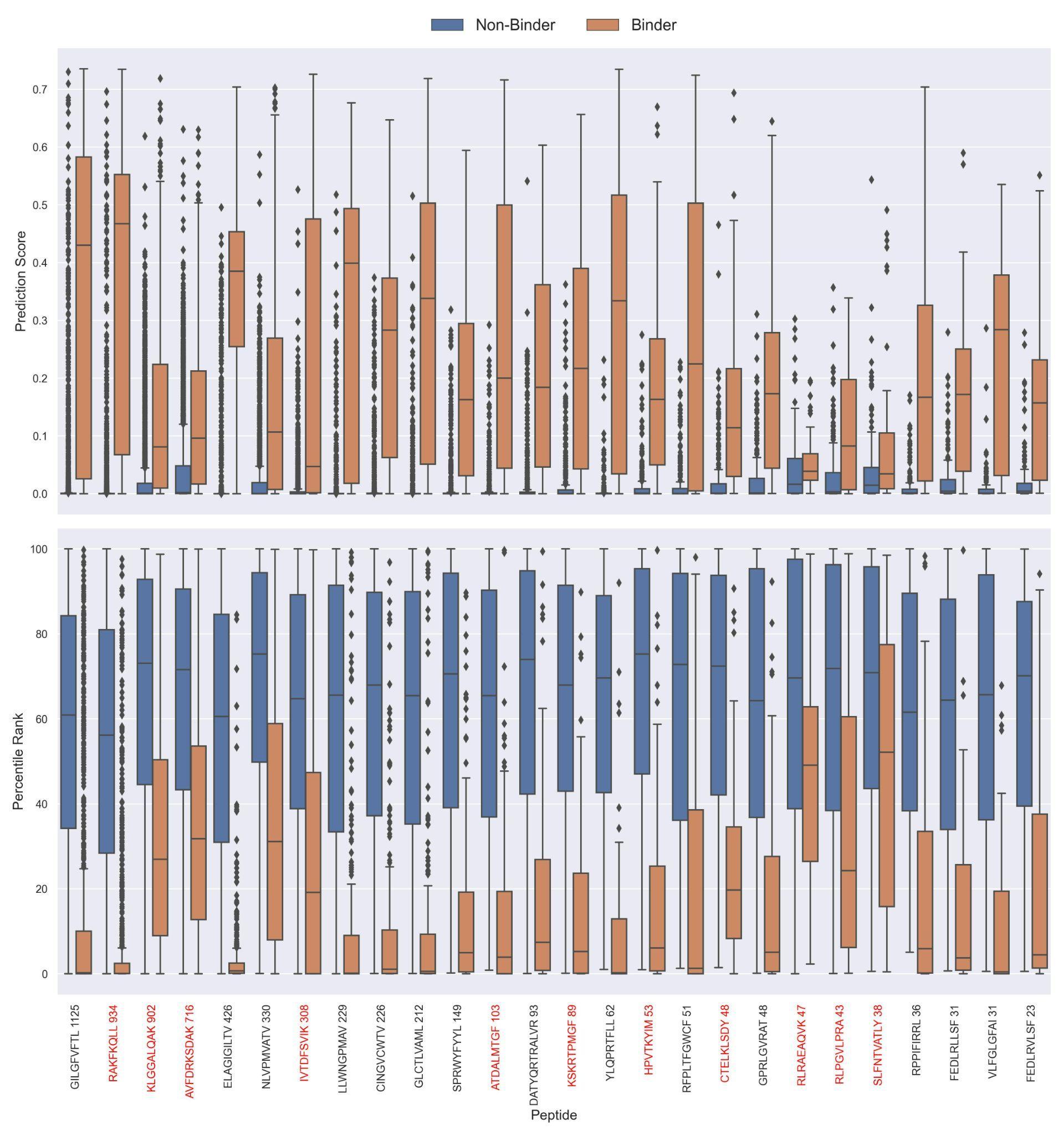

***Supplementary Figure 9*** *– Boxplot of direct prediction scores and percentile ranks per peptide of the full test dataset for the TCRbase ensemble (based on the pre-trained models). Peptides with 100% of positive observations coming from 10X sequencing are highlighted in red.*

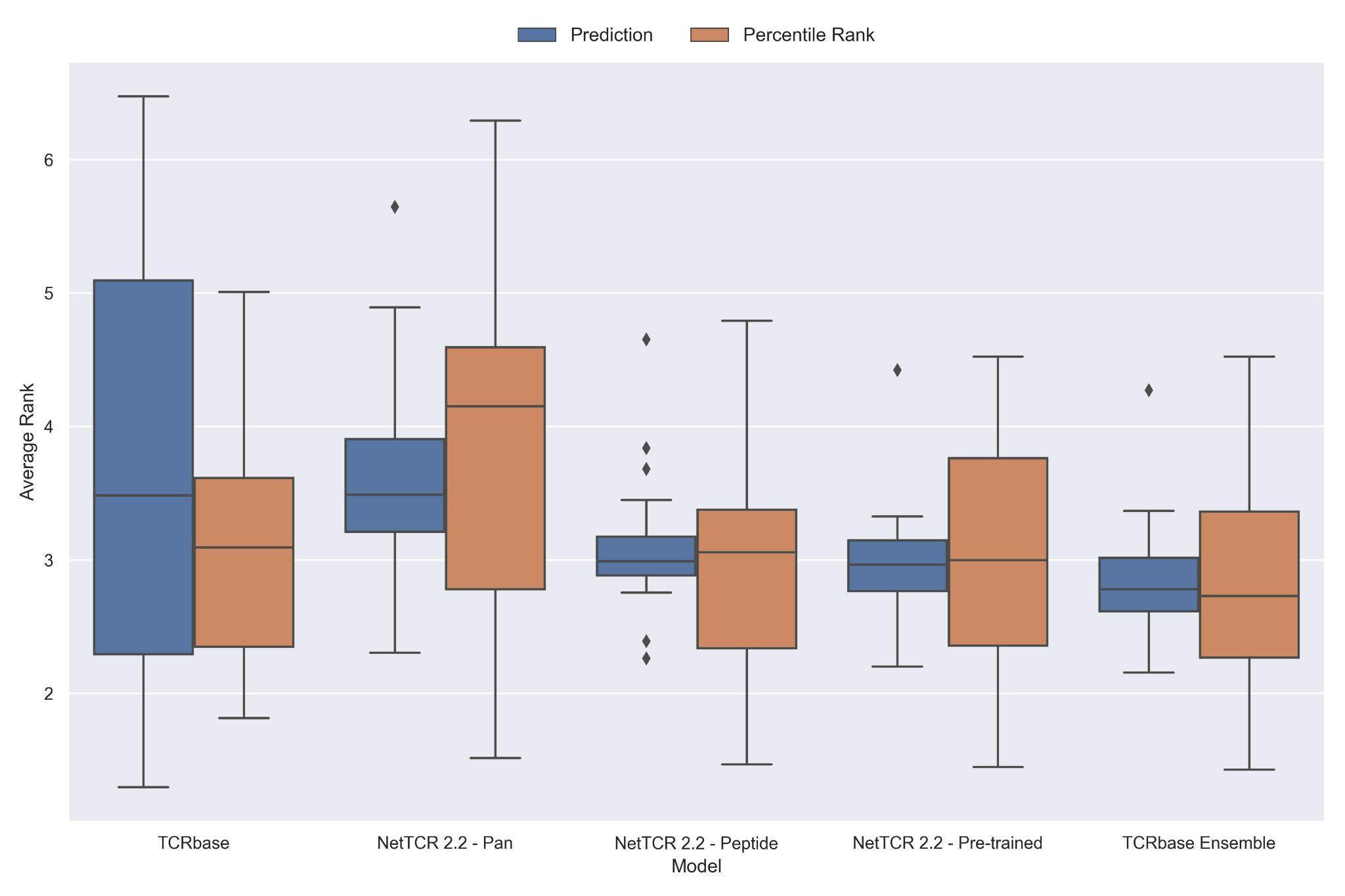

***Supplementary Figure 10*** *– Boxplot of average rank per peptide for the final updated models. The rank was evaluated on the limited dataset covering 21 peptides, i.e. excluding the peptides with low performance (KLGGALQAK, AVFDRKSDAK, NLVPMVATV, CTELKLSDY, RLRAEAQVK, RLPGVLPRA and SLFNTVATLY).*

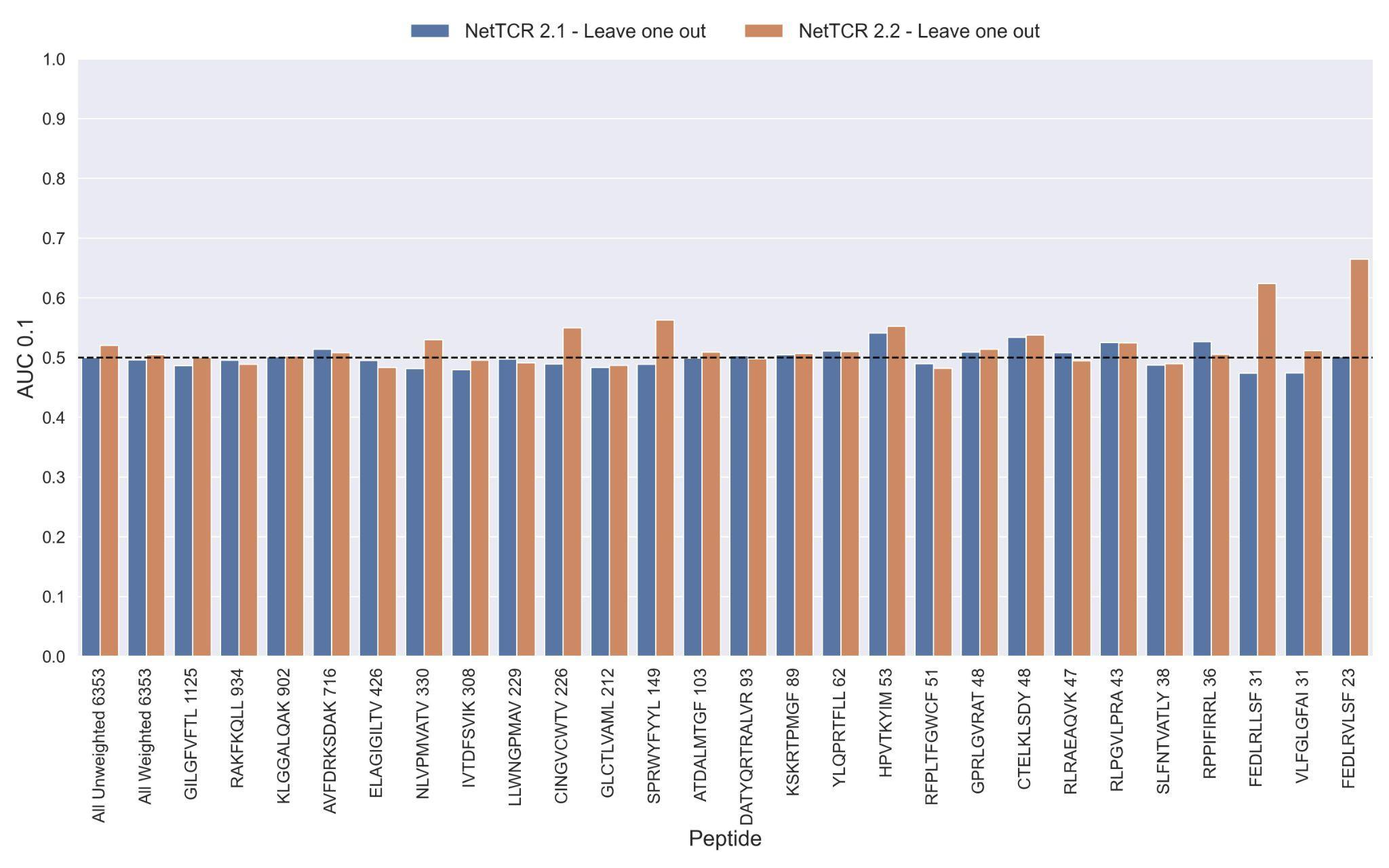

***Supplementary Figure 11*** *- Per peptide performance of the old (NetTCR 2.1) and updated (NetTCR 2.2) pan-specific CNN models trained in a leave-one-out setup. The performance was evaluated in terms of AUC on the full dataset.*

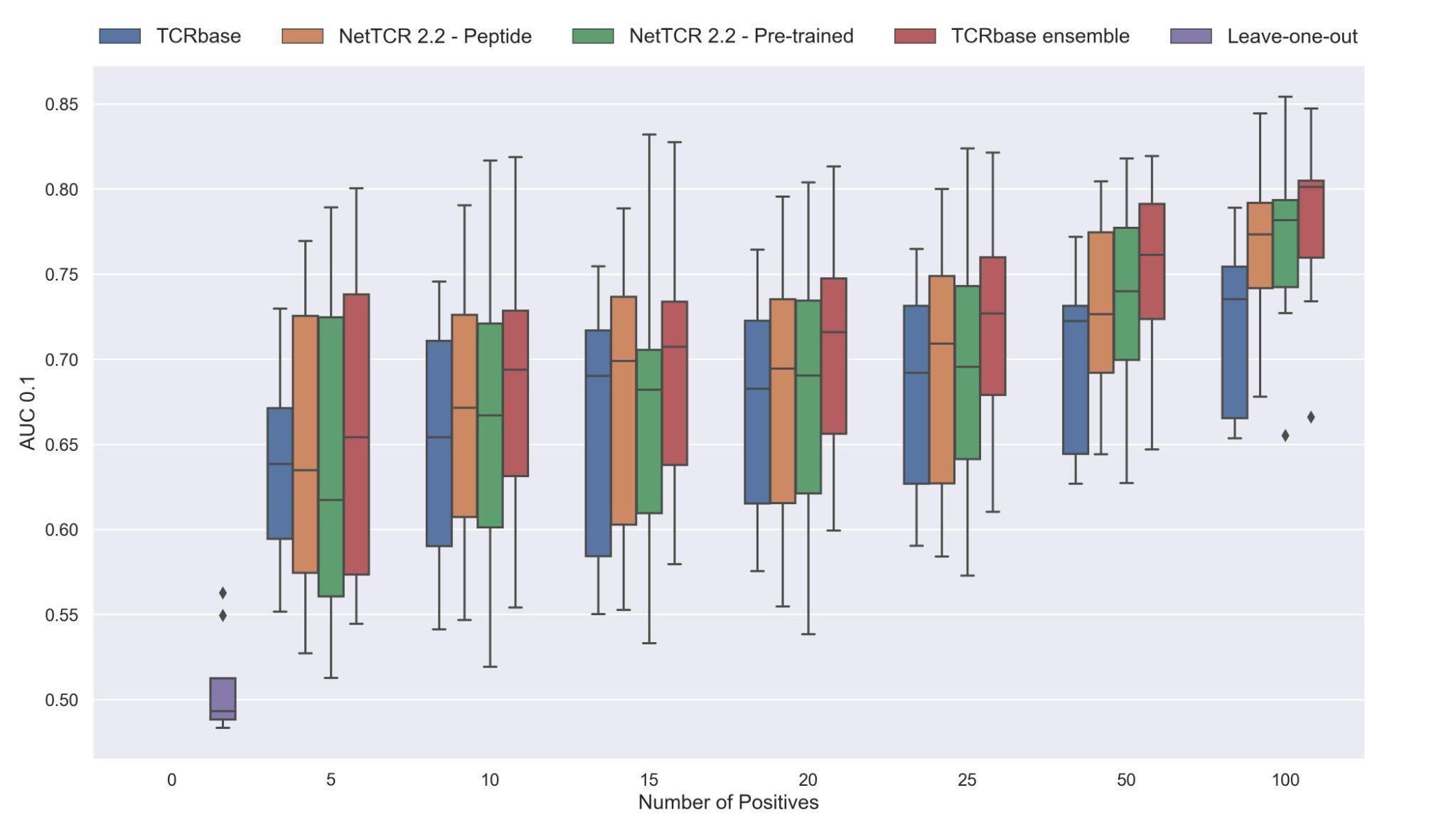

***Supplementary Figure 12*** *– Performance in terms of AUC 0.1 of various models trained on increasing amounts of data. These models were trained on the following peptides: GILGFVFTL, RAKFKQLL, ELAGIGILTV, IVTDFSVIK, LLWNGPMAV, CINGVCWTV, GLCTLVAML and SPRWYFYYL. The pre-trained models were based on the leave-one-out model, and afterwards fine-tuned and re-trained on the smaller training datasets.*

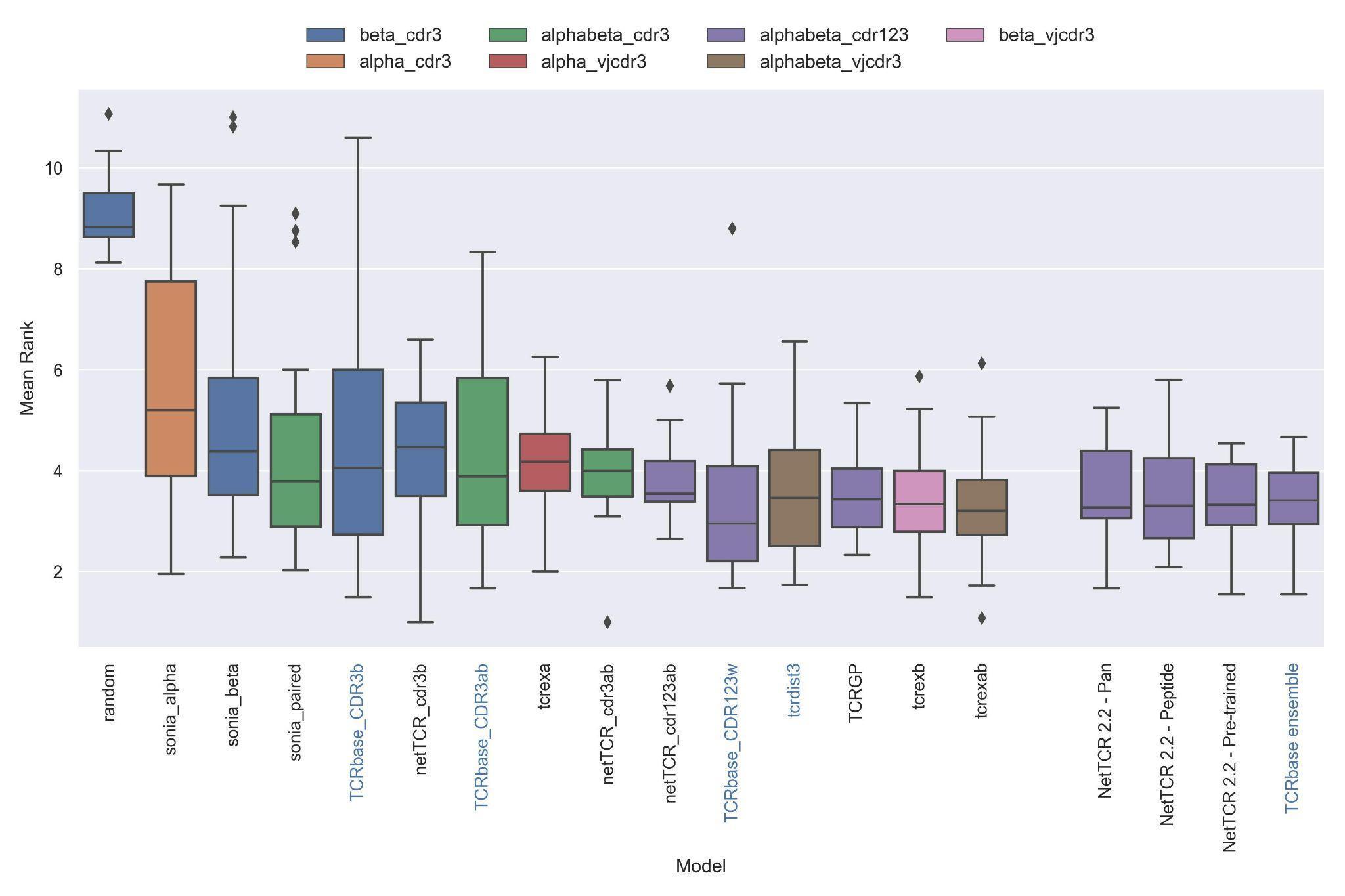

***Supplementary Figure 13*** *- Boxplot of average rank per peptide per model in the IMMREP test data, as reported in the IMMREP benchmark. The updated NetTCR 2.2 models are included to the right. The color of the bars indicates the type of input used by the model. Machine-learning models are labeled with black text, whereas distance-based models are labeled with blue text. Note that the TCRbase ensemble is a mixture between a machine-learning and distance-based model.*
